# AXOLOTL: an accurate method for detecting aberrant gene expression in rare diseases using coexpression constraints

**DOI:** 10.1101/2024.01.07.574502

**Authors:** Fei Leng, Yang Liu, Jianzhao Zhang, Yansheng Shen, Xiangfu Liu, Yi Wang, Wenjian Xu

## Abstract

**Background:** The assessment of aberrant transcription events in patients with rare diseases holds promise for significantly enhancing the prioritization of causative genes, a practice already widely employed in clinical settings to increase diagnostic accuracy. Nevertheless, the entangled correlation between genes presents a substantial challenge for accurate identification of causal genes in clinical diagnostic scenarios. Currently, none of the existing methods are capable of effectively modeling gene correlation.

**Methods:** We propose a novel unsupervised method, AXOLOTL, to identify aberrant gene expression events in an RNA expression matrix. AXOLOTL effectively addresses biological confounders by incorporating coexpression constraints.

**Results:** We demonstrated the superior performance of AXOLOTL on representative RNA-seq datasets, including those from the GTEx healthy cohort, mitochondrial disease cohort and Collagen VI-related dystrophy cohort. Furthermore, we applied AXOLOTL to real case studies and demonstrated its ability to accurately identify aberrant gene expression and facilitate the prioritization of pathogenic variants.

## Introduction

RNA-seq is an indispensable tool for identifying pathogenic DNA variants and increasing the diagnostic rate of rare disease cohorts [1–5]. For undiagnosed patients, causative genes can be further prioritized if both DNA and RNA supporting evidence are considered. Three types of extreme transcription events (abnormal gene expression, alternative splicing, and allele-specific expression) are detectable in RNA-seq data [6–10]. Aberrant gene expression is one major type of RNA evidence. Rare diseases are so diverse that cohorts usually consist of many individuals suffering from different genetic disorders. However, classical differential analysis is not suitable for accurately detecting abnormal gene expression in patients with rare diseases[1,2,11]. Specialized computational methods have been developed to detect aberrant gene expression [7,12–14].

Aberrant gene expression in an RNA expression matrix is an outlier detection problem. The existing methods measure the abnormalities of matrix elements in different ways. OUTRIDER uses a denoising autoencoder to control unwanted covariations in the gene expression profile, after which the autoencoder-normalized counts are statistically modeled with gene-specific negative binomial distributions[7]. ABEILLE uses a variational autoencoder to compute reconstruction errors, classifies normal and abnormal genes with a decision tree model in a supervised setting and quantifies anomality with an isolation forest model in an unsupervised setting [13]. The OutSingle models expression matrix with log-normal distributions, controlled for confounders with an SVD-based model, and estimates P values for genes [14]. OutPyR yields a novel Bayesian outlier score without controlling for confounders, but its performance is not superior to that of OUTRIDER [12,15]. Although recent methods can control confounders such as sequencing platform and batch, they cannot fully rule out the biological coregulation of genes [7,14]. Disentangling gene coregulations remains an unresolved challenge for detecting aberrant gene expression.

Genome-wide coregulation can be viewed as a coexpression network of many interconnected gene modules [16–18]. Inspired by that, we regard the coexpression relationship as an inherent biological constraint of an individual transcriptome. A group of coexpressing genes function like a loosely connected module, exhibiting coordinated up and down fluctuations across samples. Individual genes fluctuate within the normal variational range determined by coexpressing genes. Thus, the more a gene deviates beyond the normal variational range, the more aberrant it is in the transcription profile.

Here, we propose AXOLOTL (Aberrant gene e<u>X</u>pression as <u>O</u>ut<u>L</u>ier detecti<u>O</u>n using devia<u>T</u>ion from norma<u>L</u> coexpression), an unsupervised machine learning method to detect aberrant gene expression events in RNA expression matrix. Our work is the first to explicitly address biological confounders by introducing coexpression constraints into aberrant gene expression detection tasks. We benchmarked the AXOLOTL and existing methods using publicly available RNA-seq datasets containing aberrantly expressed genes, which included GTEx datasets of four clinically accessible tissue (CAT) samples[19,20], a mitochondrial disease (n=423) dataset of fibroblast samples[3] and a Collagen VI-related dystrophy (COL6-RD, n=36) dataset of muscle samples[21]. We also demonstrated the superior performance of the AXOLOTL model in detecting aberrant expression in our in-house dataset (n=111).

## Materials and Methods

### Real datasets

Both rare disease cohorts and healthy individual cohorts were included in this study (Figure 1A). Blood, skin and muscle tissues represent the most clinically accessible tissues (CATs) for elucidating causative genes of Mendelian diseases. GTEx subdatasets of CATs with predefined outliers[20] were used for model development and optimization. The mitochondrial disease dataset[3] and Collagen VI-related dystrophies dataset[21] were used for model performance evaluation in the rare disease cohort. In real case studies, clinical samples (ly_111 dataset) were used to highlight the practical value of aberrant gene expression.

**Figure 1.**
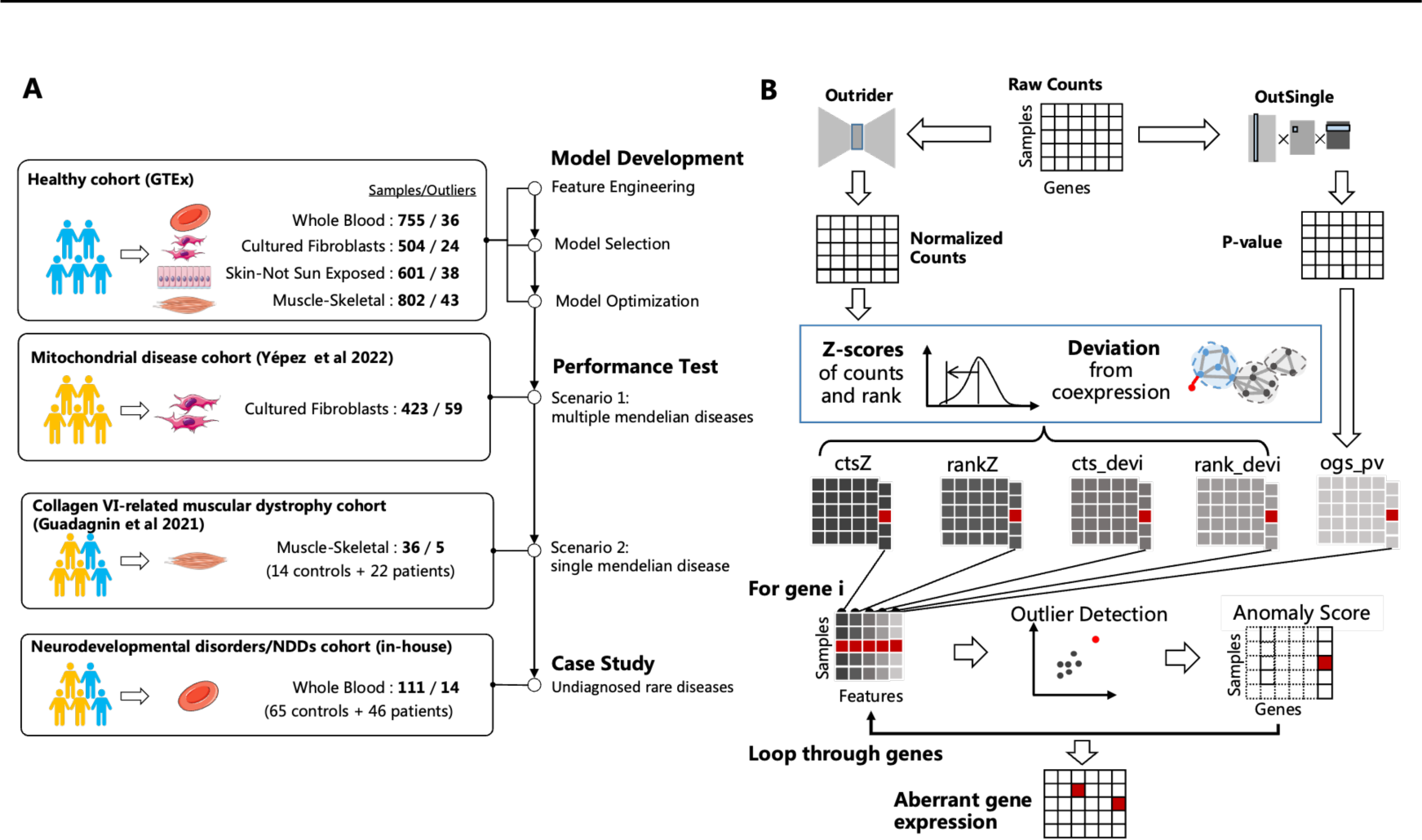
Workflow and method overview of AXOLOTL. **(A)** Datasets. Schematic of cohorts, tissues, RNA-seq datasets and their roles in different stages of this study. Human shaped icons represent controls (blue) and rare disease patients (yellow). For each dataset, the samples represent the number of biological specimen and the outliers represent the number of sample-gene pairs with aberrant gene expression events. **(B)** AXOLOTL architecture. Input is an RNA-seq read count matrix. Rows represent samples; columns represent genes. Gene read counts are processed into our customized features, which characterize how far each gene-sample pair deviate from its normal range. Genes correspond to columns of those features; the same column of five features are reorganized together as feature table for a gene. Unsupervised outlier detection model fit on the feature table of one gene at a time. Model outputs anomaly score. Aberrant expression events at specific sample-gene pairs are highlighted in red.

### GTEx cohort: blood_755, skin_601, fib_504 and muscle_802

RNA-seq read counts (v8) were downloaded from the GTEx portal. The dataset consisted of 17,382 transcription profile data from 54 healthy tissues of 948 donors. Each profile contains expression levels of >10,000 genes. Whole-blood, skin-not sun exposed, cultured fibroblast and muscle-skeletal samples were extracted as four GTEx datasets: blood_755, skin_601, fib_504 and muscle_802 (Figure 1A, Supplemental Methods). Among the global multi-tissue outliers [20], we defined aberrant expression with an arbitrary cutoff at Z score >9 or <-7 (Supplemental Table S1∼4). There was approximately 1 aberrantly expressed gene per 20 samples.

### Rare disease cohort 1 - pfib_423

Pfib_423, the largest publicly available rare disease RNA-seq cohort dataset to date, is a dataset of skin-derived fibroblasts from 423 patients (including the Kremer dataset[1]) suffering from mitochondrial disease or other Mendelian diseases[3]. The read count table was retrieved from Zenodo 4646823 and 4646827. We compiled 59 positive aberrant expression events (Supplemental Table S5). All the positives were reported in the aberrant expression analysis and verified by WES/WGS in original publication.

### Rare disease cohort 2 – pmuscle_36

Pmuscle_36 is an RNA-seq dataset of skeletal muscle biopsies from 22 COL6-RD patients and 14 age-matched controls [21]. COL6-RD is a form of congenital muscular dystrophy characterized by muscle weakness and joint defects. The molecular mechanism of COL6-RD involves loss of function or dysfunction of COL6 microfibrils in the muscle extracellular matrix. COL6-RD can be caused by pathogenic variants in collagen VI genes (COL6A1, COL6A2, and COL6A3). Pathogenic variants in COL6A genes were found in 22 patients. Immunostaining assays indicated that COL6A proteins were aberrantly expressed at low levels in 5 patients and were mislocalized in the other 17 patients. We defined the COL6A genes of these 5 patients as expression outliers (Supplemental Table S6). Gene read counts were produced using our in-house pipeline described previously [22] from the RNA-seq raw dataset GSE103975.

### Rare disease cohort 3 – ly_111

This study was approved by the Institutional Review Board of the Capital Institute of Pediatrics (HERLLM2023060, SHERLLM2023061). Informed consent was obtained from all the subjects participating in the study. The research conformed to the principles of the Declaration of Helsinki. 65 controls and 46 probands suffering from inherited disorders of the nervous system were retrospectively enrolled. The samples were accepted and evaluated by a genetic laboratory (Supplemental Material). All probands had DNA variants predicted to cause RNA alternative splicing (AS) of clinically suspected pathogenic genes (Supplemental Table 12). The control group consisted of 65 miscellaneous samples, including (a) unaffected parental samples of some probands and (b) probands and parents not suitable for blood specimen-based aberrant expression analysis. All the samples were subjected to blood RNA-seq and sequenced on the same platform. Gene read counts were generated with an in-house pipeline. To validate the splicing effect of the candidate variants, alternative splicing events proximal to the variants were manually inspected. Variants validated by aberrant splicing were considered pathogenic. Variants that were not validated were identified as VUS. After RNA-seq alternative splicing analysis, 30 cases were resolved, and 16 cases were undiagnosed.

See the Supplemental Material for more details about dataset preprocessing.

### Semi-synthetic dataset for robustness tests

*Pfib_423* is the largest rare disease RNA-seq dataset in this work. We used *Pfib_423* to analyze the robustness of AXOLOTL. Robustness refers to the ability of a method to consistently outperform other methods across multiple dataset settings, including varying sample sizes and percentages of patients with aberrant gene expression. We tested a total of 15 combinations of sample sizes [100, 150, 200, 250, 300] and percentages of positive samples [0.05, 0.10, 0.15]. The raw pfib_423 dataset was randomly subsampled for the desired combination settings (e.g., sample size = 150 and percentage = 0.10) to obtain the simulated dataset. The performance of each combination setting was averaged across 100 simulations with different random seeds.

### AXOLOTL method

An overview of the AXOLOTL method is shown in Fig. 1B. We next describe the implementation details of feature construction and its machine learning model.

### New features based on coexpression deviation

To construct a better model for detecting aberrant gene expression, the deviation of gene expression from the coexpression constraint was measured by newly customized features. The gene read count matrix !_!_represents “ genes (in row) and # samples (in column). Each element of matrix !_!_represents a gene-sample pair !_!_(%, ’). In this section, we define five customized features (*ct*s_z, *rank_z*, *cts_dev*, *rank_dev*, and *ogs_pv*) and explain why they quantitatively characterize how far each gene-sample pair deviates from its normal coexpression. First, the gene read counts were preprocessed through OUTRIDER autoencoder normalization.

The encoder learns from low-dimensional encodings from raw expression profiles, and the decoder regenerates normalized gene expression profiles from latent encodings. The confounding factors (batch effect, technical variation and sequencing depth) in !_!_ were controlled by a denoising autoencoder to create a normalized gene expression table !. The gene expression levels of ! are more comparable across samples than are those of !_!_. The optimal latent encoding dimension of the autoencoder is searched within integers ranging from 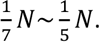.

Second, the z scores-based features were computed. The expression matrix ! is standardized across all samples to construct the z score matrix *ct*s_z. Then, within every column of !, the genes were ranked in descending order of expression level. Gene rankings serve as “relative” expression levels. Changes in the rankings are usually subtle and less affected by overdispersion in gene read counts[11]. For each gene, standardized rankings of the same gene across all samples were used to obtain the z score matrix *rank_z*.

Third, coexpression partners of genes were identified. Pearson’s correlation coefficient * was computed for all genes from the normalized expression table !. Assume * is an ” × “ matrix, where *(%, %′) represents the coexpressing strength of genes % and %′. The closer *(%, %′) is to 1, the more strongly two genes are linearly correlated. We defined the genes %^%^ with the 2% largest *(%, %′) as coexpressing genes % %^∗^.

Fourth, we measured the deviation of gene expression from the coexpression constraint using the average z score difference between gene % and its coexpressing partners %^∗^ . The two deviation features of the gene‒sample pair (%, ’) are computed based on the corresponding z score features:

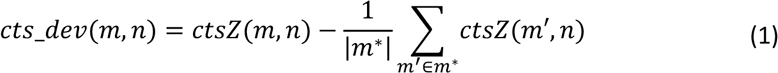

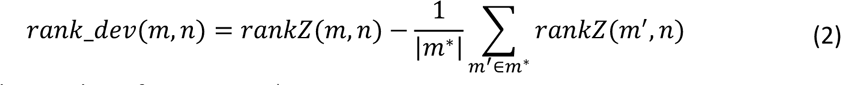

where |*m*^∗^| is the number of genes in *m*^∗^.

Finally, we adopt the =-value statistic of the OutSingle method as one additional feature. The =-value matrix is negative-logarithmically transformed and denoted as feature matrix *ogs_pv*. This feature aims to quantify whether the gene expression level follows a negative binomial distribution from a statistical perspective [14].

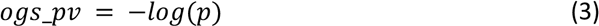

### Outlier detection model

The outlier detection model is trained for each gene. For the model of gene %, the input feature matrix is # × 5, where samples is in rows and features is in columns. The model detects which samples have aberrant expression of genes %. The model assigns anomaly scores for all genes in a sample. Specifically, an unsupervised machine learning model called the local outlier factor (LOF) is used to detect aberrant gene expression in AXOLOTLs. The LOF estimates the local density of a sample by considering the distance between its k-nearest neighbors. A lower local density indicates an outlier sample with respect to the others.

In the model explanation section, two outlier detection models were chosen to prove the suitability of LOF in AXOLOTL (Fig. 3C). The first model is Isolation Forest (IF). It builds a recursive partitioning tree structure by randomly selecting one feature and then randomly splitting by the selected feature. The number of splits needed to reach a sample from the root node is the path length of the sample. Extreme short average path suggests the sample may be an outlier. The second model is One Class Support Vector Machine (OC-SVM). The OC-SVM assumes that normal samples are mainly from one class with smooth boundaries determined using input data. Samples far away from the normal class boundary are outliers.

### Performance evaluation

Our optimal model was employed for comparative analysis with representative baseline models, which are OUTRIDER [7], ABEILLE [13] and OutSingle [14]. Their performance on detecting aberrant gene expression was tested on the labeled GTEx cohort and rare disease cohorts of different tissues. We collected the outputs of the gene-specific models and reorganized them in the same shape as the input matrix. The gene rankings in the sample were more useful than the raw anomality score. Outliers called by a method need to be checked manually by doctors or genetic counselors. It is desirable that aberrantly expressed genes are among the top-ranking genes in this sample. The Precision-Recall (PR) curve and the area under the PR curve (AUPRC) are used to evaluate the performance of the models. The AUPRC, a metric for a class imbalanced dataset, is particularly useful for us because it places significant emphasis on the model’s ability to correctly identify positives. Model performance was also evaluated with the top- < hit recall metric, i.e., the proportion of disease-relevant known outlier genes among the top < genes per proband sample. To achieve this, anomality scores of genes were ranked within each sample to prioritize the most aberrant genes and compute recall at predefined top-< hits.

### Feature attribution and Ablation analysis

Feature attributions are analyzed by the model-agnostic explanation methods SHAP (SHapley Additive exPlanations, [23]) and LIME (Local Interpretable Model-Agnostic Explanations, [24]). Feature attributions indicate how much each feature contributed to our model’s output for samples of a given dataset. The two rare disease datasets pmuscle_36 and pfib_423 were used. For pfib_423, gene-specific models of disease-causative genes (52 unique genes; Supplemental Table S1) were included. For pmuscle_36, the gene-specific models of disease-causing genes (COL6A1, COL6A2, and COL6A3) of COL6-RD, causative genes of other phenotypically similar diseases (COL6A6, CAPN3, DYSF, TTN, TCAP and LMNA) and 100 random genes were included.

Ablation experiments are carried out to test the components of the AXOLOTL. On the feature side, one or two features were removed to generate ablated feature sets. On the model side, the outlier detection model was changed from LOF to IF or OC-SVM. We compared the performances of these candidate models with those of AXOLOTL on multiple datasets.

### Implementation

The LOF, IF and OC-SVM models were implemented using scikit-learn v0.21 with default parameters unless otherwise specified: neighbors.LocalOutlierFactor (n_neighbors=20), ensemble. IsolationForest (n_estimators=20, max_samples =1.0), and svm. OneClassSVM (gamma=’auto’). Pearson’s correlation coefficient was calculated with scipy.stats from scipy v1.3 [35]. Feature importance analysis of SHAP and LIME was performed with scikit-explain v0.1.2. The plots were drawn using Matplotlib v3.1 [37] and Seaborn [25] in Python-3.7. OUTRIDER v1.12.0 and ABEILLE v1.0.0 were implemented in R packages running on R-4.1.1. RNA AS was visualized using an Integrative Genomics Viewer [26].

## Results

### Overview of the AXOLOTL method

AXOLOTL detects aberrant gene expression using RNA-seq read count data (Fig. 1B). The input of AXOLOTL is solely an RNA-seq read count *N_gene*×*N_sample* matrix. The output of AXOLOTL is an anomaly score matrix with the same shape as the input. Then, a set of customized features were constructed to measure the deviation of gene expression from the coexpression constraint. Using these newly designed features, an unsupervised outlier detection model was built to detect rare aberrant gene expression events.

Aberrant gene expression identification of each gene is an anomaly detection task. The workflow consists of three steps. First, we normalize the gene expression matrix by OUTRIDER’s autoencoder component. The normalized gene expression level was more comparable across samples than was the raw read count. Second, we constructed five features capturing the deviation of a gene from its normal expression range. By comparing one gene across many samples, standardized z-score features *cts_z* and *rank_z* are used to quantify how aberrant it is in each sample. Furthermore, for AXOLOTL, Pearson’s correlation coefficient (*) was used to measure the strength of coexpression between genes. For one gene, other genes among the top 2% largest * were defined as coexpression partners. The coexpression partner relationships serve as constraints on gene expression variations in normal samples. The features *cts_dev* and *rank_dev* measure how much a gene deviates from its coexpression partners in the same profile. The last feature, *ogs_pv,* is the negative log-transformed p value outputted by OutSingle. Third, for each gene, we fit an unsupervised outlier detection model using a gene-specific matrix of *N_sample* rows and 5 columns. Genes were looped through iteratively to detect aberrant gene expression.

### AXOLOTL detects healthy individuals’ gene expression outliers

Healthy individuals in the GTEx database exhibit extreme outlier gene expression due to rare genetic variations[20]. Since gene expression outliers are caused by rare genetic variations in the healthy population as well as rare disease patients, the gene expression dataset of GTEx donors is suitable for unbiased benchmarking of aberrant gene expression detection methods. From the GTEx database, we collected expression profiles of CATs to prepare 4 datasets, namely, blood_755, skin_601, fib_504 and muscle_802, which contained 36, 24, 43 and 38 known gene expression outliers, respectively (Fig. 1A). We compared the performance of the AXOLOTL with that of three baseline models in detecting outliers in healthy individuals.

The shapes of the PR curves produced by all the methods were quite similar for one dataset (Figure 2A). We observed the relative superiority of AXOLOTL over other methods across datasets. Specifically, the AXOLOTL performed the best on the most accessible blood tissue, indicating its clinical advantages. AXOLOTL also has slightly higher precision in the recall range of 0.5∼0.6 in the other three datasets. A comparison of AUPRC further confirmed the above results (Fig. 2A). Next, the recall of top-k hits is used to quantify how well a method prioritizes real outliers as the top-ranking genes per sample. We observed that the performance of AXOLOTL was highly competitive in terms of top-k hit recall curve (Figure 2B). In summary, AXOLOTL outperformed the other methods in detecting gene expression outliers in healthy individuals.

**Figure 2.**
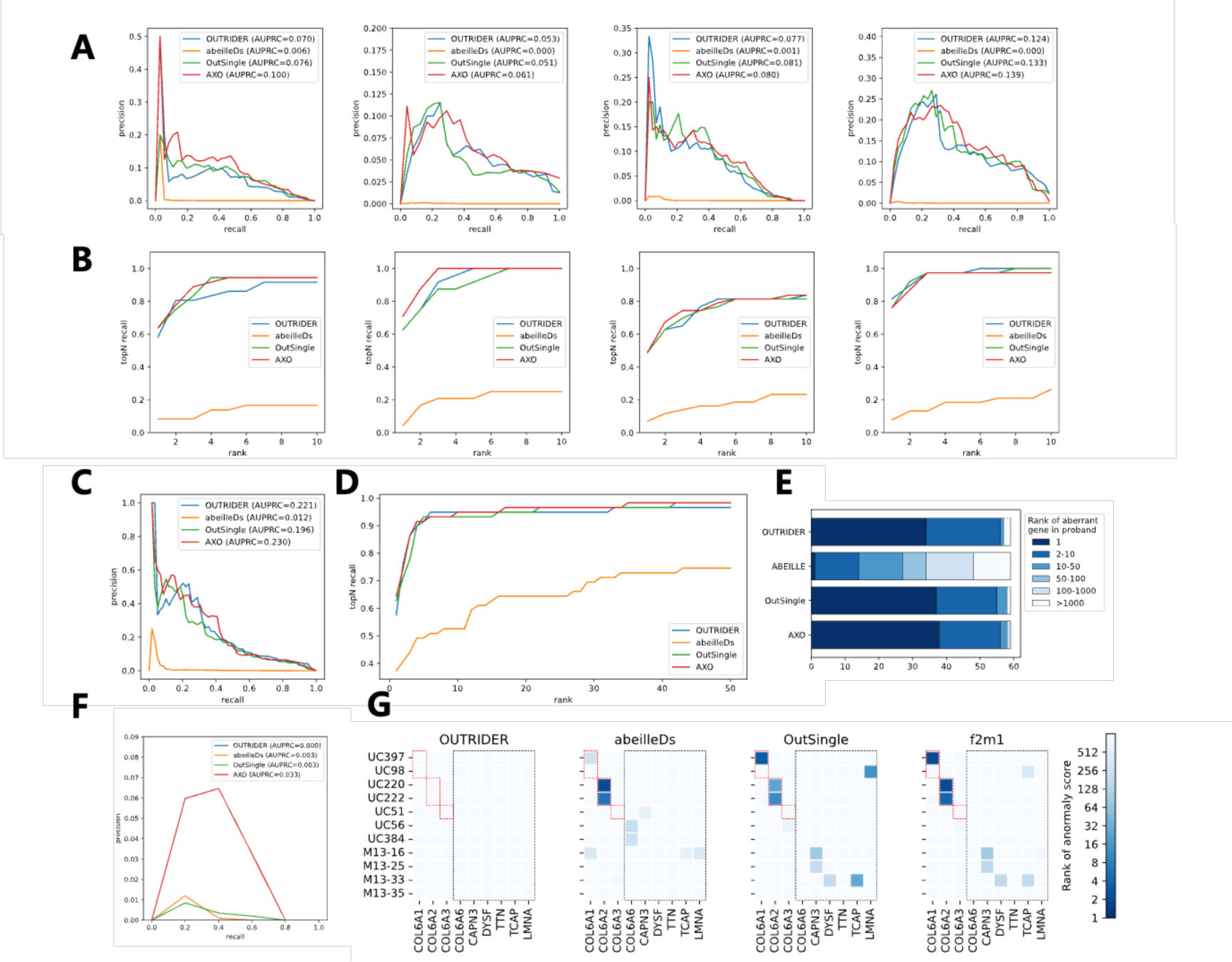
Performance comparison of AXOLOTL with other methods on public datasets. Performance on GTEx datasets of 4 clinically accessible tissues from healthy donors: **(A)** Precision Recall (PR) curve and Area Under the PR curve (AUPRC) and **(B)** Top-k hit recall curve, where k is 1∼10. Performance on pfib_423 dataset of skin-derived fibroblasts: **(C)** PR curve and AUPRC, **(D)** Top-k hit recall curve, where k is in the range of 1∼50 and **(E)** ranking distribution of aberrant gene in proband. Performance on pmuscle_36 dataset of skeletal muscle biopsies: **(F)** PR curve and AUPRC and **(G)** rankings of COL6-RD causative genes (COL6A1, COL6A2, COL6A3) and six other disease genes (COL6A6, CAPN3, DYSF, TTN, TCAP and LMNA) with similar phenotype. Color of squares represent log2 scaled of rankings. Red dashed edges indicate the true positives. Black dashed edges indicate false positives genes.

**Figure 3.**
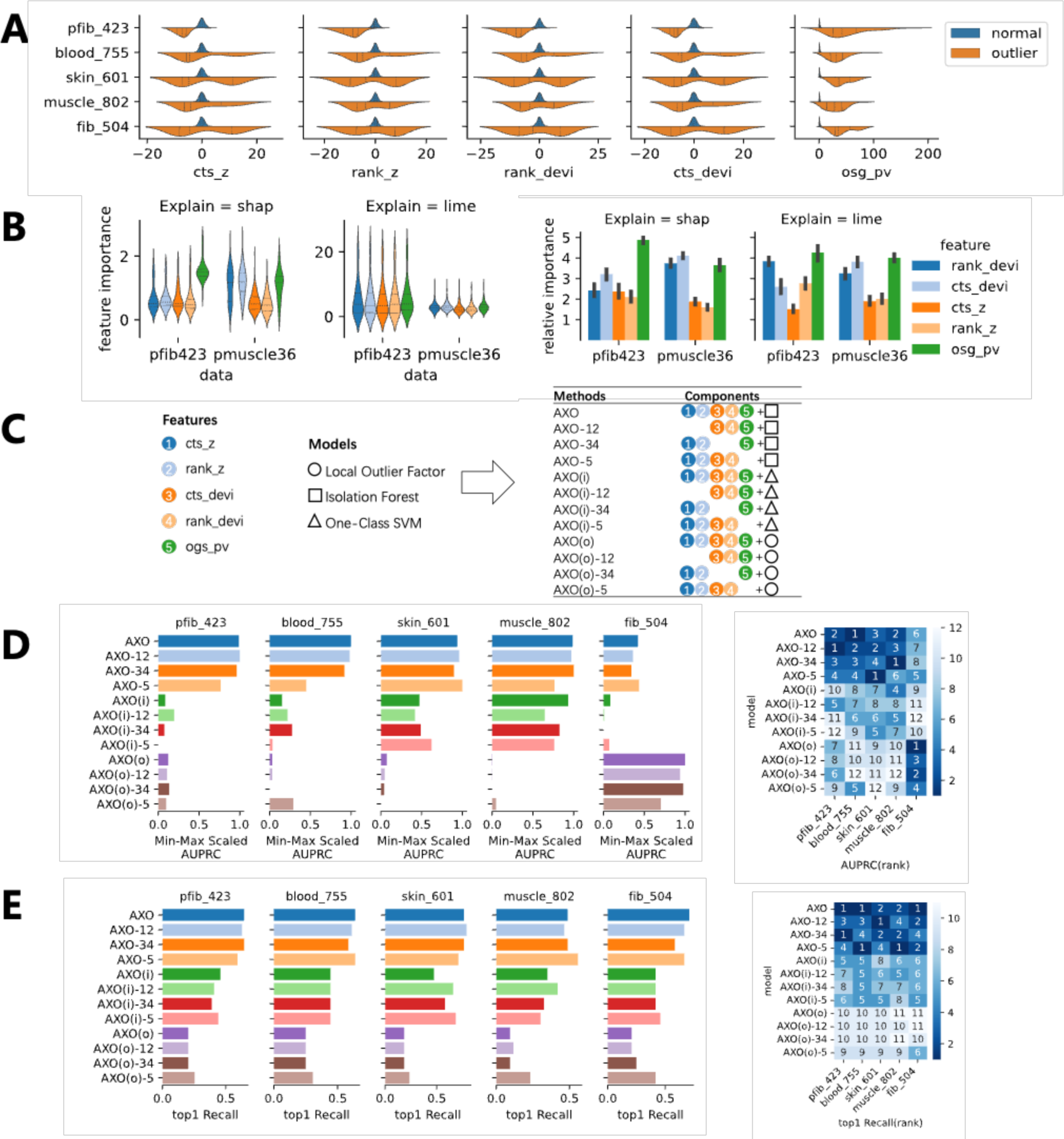
Model explanation by feature contribution and ablation analysis. **(A)** Value distributions of each feature on GTEx and pfib_423 datasets. Aberrantly expressed gene- sample pairs and a random subset of other gene-sample pairs are visualized. **(B)** Feature attributions on rare disease datasets pmuscle_36 and pfib_423 are analyzed by SHAP and LIME. Left panel: violin plots show the feature attribution of gene-specific outlier detection models of disease causative genes, non-causative genes and a group of random genes. Right panel: bar plots show the relative importance of features on each of those outlier detection models. **(C)** schematic of five features, three candidate outlier detection models, and the experimental setup of ablation experiments. Color filled circles represent features. Three blank shapes represent models. AXOLOTL is abbreviated as AXO. Other 11 settings are combinations of feature ablation and model replacement on AXO method. **(D)** Area Under the PR curve (AUPRC) of candidate combinations of feature sets and models in GTEx datasets and pfib_423. Left panel is Min-Max scaled AUPRC. Right panel is rankings of AUPRC. **(E)** Top-1 hit recall of candidate combinations of feature sets and models in GTEx datasets and pfib_423. Left panel is top-1 hit recall. Right panel is rankings of top-1 hit recall.

### AXOLOTL outperforms existing methods on rare disease cohorts

We next evaluated the diagnostic performance of the AXOLOTL in two rare disease cohorts.

The pfib_423 dataset is a multidisease cohort of skin-derived fibroblasts from patients with mitochondrial disease or other Mendelian diseases [3]. pfib_423 is an extension of the widely used Kremer dataset [1]. A total of 59 patients were solved with causative genes confirmed by aberrant gene expression analysis. Here, these 59 sample–gene pairs are labeled as positive points in the pfib_423 dataset. Because the pfib_423 dataset has 3 times more total samples and 7 times more positives than does the Kremer dataset, a model performance comparison of pfib_423 could be more convincing. The PR curves and top-k hit recall curves of different methods are shown in Fig. 2C-D. It seemed that four methods tend to coincide on most of positive events (Supplemental Figure 1). In detail, AXOLOTL has the highest AUPRC. AXOLOTL more likely recovered true outliers as a top-1 aberrant gene in probands (Figure 2E and Supplemental Table 5). OUTRIDER is the second-best method in terms of AUPRC; OutSingle is the second-best in terms of the top-1 recall. Considering all the outliers discovered by OUTRIDER, the relative advantage of AXOLOTL over OUTRIDER indicates that AXOLOTL successfully raises the upper limit of outlier detection performance. These results prove the superiority of AXOLOTL over other baseline methods for detecting aberrant gene expression in a large cohort.

A rare disease cohort such as pfib_423 [1–5] is idealistic but difficult to construct because of the low prevalence of rare diseases in population. A small cohort of a few tens of patients’ RNA-seq data is more often available. The performance of AXOLOTL on such a small dataset reflects its reliability in diagnostic scenarios. One example is the pmuscle_36 dataset, which includes 22 patients with COL6-RDs and 14 controls [21]. Pathogenic variants of COL6A1, COL6A2 or COL6A3 genes were found and validated by collagen immunostaining in all COL6-RDs. COL6 protein and mRNA levels were reduced in 5 of the 22 patients. Known pathogenic genes with reduced mRNA levels were labeled 5 true outliers in the pmuscle_36 dataset. We benchmarked AXOLOTL and other baseline methods on pmuscle_36 (Supplemental Table 6). We observed that AXOLOTL outperformed other methods in terms of PR curve and AUPRC (Figure 2F). At the recall rate of 0.4, the precision of the AXOLOTL (0.063) was 6 times higher than that of other models (<0.010). AXOLOTL detected more known positives as the top 10 most aberrantly expressed genes in patients (Fig. 2G). In contrast, OUTRIDER detected no outliers, ABEILLE detected fewer true positives, and OutSingle assigned higher rankings to false positives in non-COL6-RD genes and lower rankings to true positives (Fig. 2G). All methods failed to detect another two outliers, i.e., COL6A1 of UC98 and COL6A2 of UC51 (Supplemental Table 6).

Taken together, AXOLOTL outperforms other baseline methods in detecting expression outliers in representative rare disease datasets. Notably, AXOLOTOL exhibited a substantial advantage when applied to smaller cohorts.

### Model explanation and feature ablation

To facilitate a comprehensive understanding of the relative importance of the five customized features in our optimal model, we next interpret the strength of AXOLOTL via feature attribution-based explanation. The feature attributions reveal relative importance of the features in explaining the output of AXOLOTL. For each feature, aberrantly expressed gene-sample pairs had larger absolute values than did normal pairs (Figure 3A and Supplemental Table 7). We analyzed feature importance using model-agnostic methods, game theory-based SHAP and linear model-based LIME. We performed SHAP and LIME analyses on the rare disease datasets pmuscle36 and pfib_423 (Supplemental Table 8). The SHAP and LIME attributes had large ranges of variation across the selected sample–gene pairs (Fig. 3B left panel). However, the relative rankings of features’ attribution on each sample–gene pair were quite stable (Fig. 3B right panel). The features, sorted from highest to lowest importance according to the SHAP and LIME values, are the *p* value statistic (*ogs_pv*), coexpression deviations (*cts_dev* and *rank_dev*) and z-scores (*ct*s_z and *rank_z*). Interestingly, both coexpression deviations and *p-*value were important for pmuscle36. These results suggest that coexpression features provide useful supporting information in small cohorts. We then implemented a total of 11 candidate combinations of feature subsets and machine learning models (Fig. 3C). Feature subsets were generated by ablating one or two features. The outlier detection model included LOF, IF, and OC-SVM. We systemically compare the performances of the above candidates with those of AXOLOTL. When LOF was changed to another outlier detection model or when any features were ablated, the overall performance of AXOLOTL was compromised in terms of AUPRC and top-1 recall (Fig. 3D-E and Supplemental Table 9). Based on these findings, we concluded that AXOLOTL is the optimal choice for its intended purpose.

### AXOLOTL is robust across various dataset compositions

To further explore the robustness of our AXOLOTL method, we simulated a series of datasets from pfib_423 to represent the different conditions (Methods). Because both rare disease cohorts have 14% of known positive samples (59/423 for pfib_423 and 5/36 for pmuscle36), a lower percentage of positive samples and a smaller cohort size were considered two factors impacting the performance of the methods. Therefore, we benchmarked AXOLOTL at 5%∼15% of positive samples and the cohort sizes ranged from 100 to 300 (Supplemental Table 10-11).

First, we analyzed the performance robustness with AUPRC (Supplemental Table 10). The AUPRC of AXOLOTL and baseline methods also positively correlated with the percentage of positive samples (Fig. 4A). When 5% of the samples were positive, the AUPRC increased from a cohort size of 100 to 150 and remained constant above 150. With 10% and 15% positive samples, AXOLOTL was the best overall, except for the cohort size of 100. Second, we analyzed the performance robustness with respect to the top-10 hits recall. AXOLOTL is superior to other methods across cohort sizes ranging from 100 to 250. AXOLOTL and baseline methods were not significantly affected by the percentage of positive samples (Fig. 4B). OUTRIDER is still a strong competitor of AXOLOTL in some cases. For example, under the condition of 5% positive samples or a cohort size of 100, OUTRIDER has a better AUPRC, but AXOLOTL has a better top-10 recall. To elucidate the comparison, we examined the top 1∼50 recall curves for the cohort size of 100 and 5%∼15% positive samples (Supplemental Table 11). AXOLOTL apparently identified more true outliers than did OUTRIDER (Figure 4C). These results suggest that AXOLOTL may have different strengths than OUTRIDER. We concluded that AXOLOTL is robust to various cohort sizes and percentages of positive samples.

**Figure 4.**
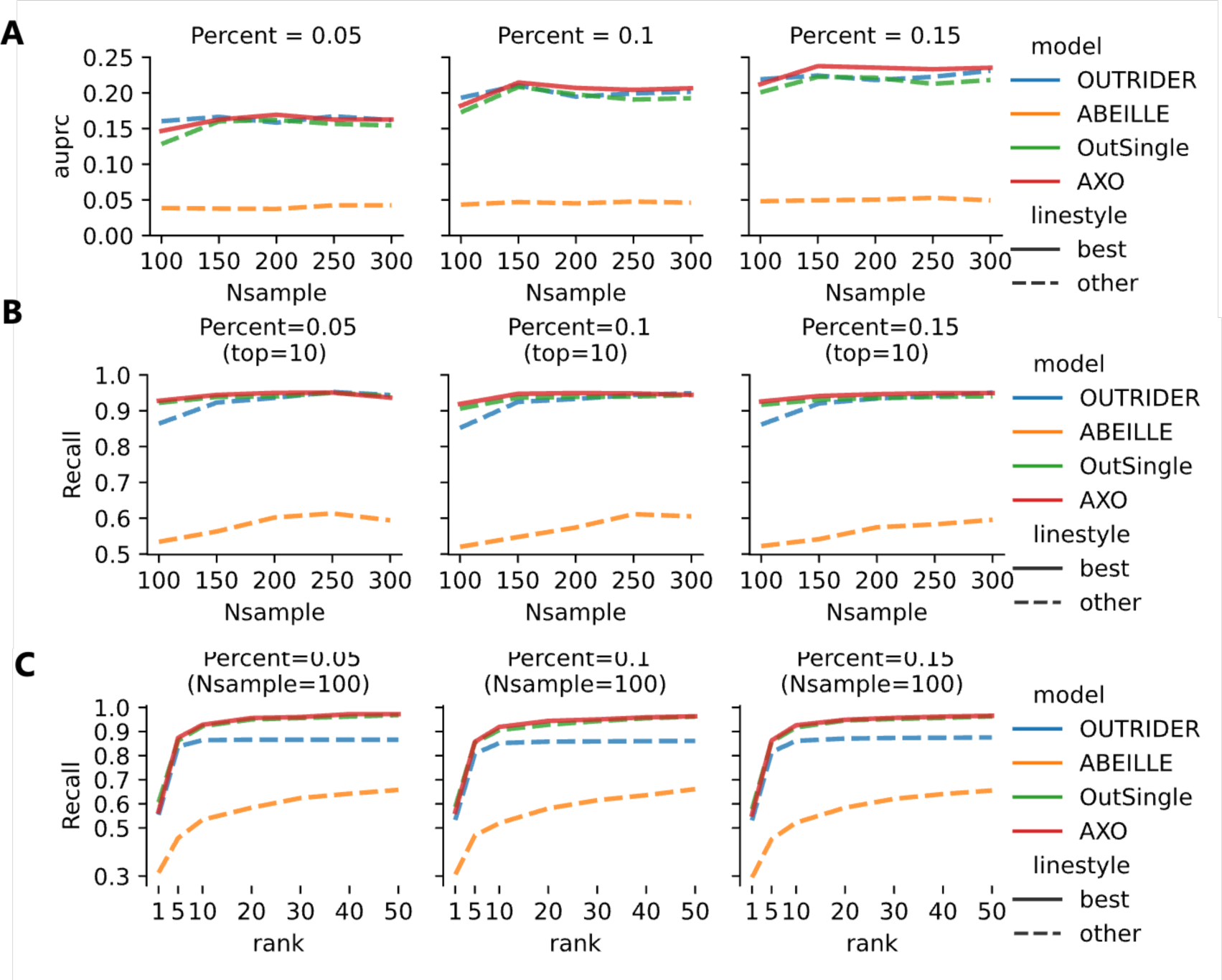
Robustness analysis of AXOLOTL at various cohort sizes and percentages of positive samples. Datasets of specific cohort sizes are random subsampled from pfib_423 dataset. 100 simulations are conducted at each cohort size and positive percentage. Average value of 100 simulations was computed for every condition. **(A)** Area Under the PR curve (AUPRC) at cohort size 100∼300 at 5%, 10% and 15% positive samples. **(B)** Top-10 hit recall of at cohort size 100∼300 at 5%, 10% and 15% positive samples. **(C)** Top-1∼50 recall at cohort size 100 at 5%, 10% and 15% positive samples.

### Real case study

We have demonstrated that AXOLOTL outperforms baseline methods in the analysis of fibroblast-based pfib_423 and muscle-based pmuscle_36 datasets. We next aimed to explore its potential in facilitating the filtering and interpretation of results derived from a blood-based RNA-seq cohort of rare disease samples.

We applied the AXOLOTL method to the in-house ly_111 dataset to identify aberrant gene expression using blood RNA-seq. The dataset is composed of 65 controls and 46 probands mainly with neural developmental disorders. Most probands had phenotypes associated with causal DNA variants predicted to affect splicing by SpliceAI (Supplemental Methods and Supplemental Table 13). We first manually checked whether aberrant splicing events existed near these variants. Considering both the evidence of DNA-level support and the presence of RNA alternative splicing (AS) near pathogenic variants, 30 cases were resolved, and 16 cases remained undiagnosed. Next, we used AXOLOTL to identify aberrant expression events independently and searched for supporting evidence for both the diagnosed and undiagnosed cases.

In 9 out of the 30 diagnosed cases, AXOLOTL successfully validated aberrant expression of the pathogenic genes (Figure 5A). AXOLOTL detected aberrant expression of genes with VUS variants in 4 out of the 16 undiagnosed cases, indicating their potential pathogenicity (Fig. 5A). Overall, aberrant gene expression was found in 28.3% of probands with suspected candidate variants. The aberrant genes of these cases were ranked higher by AXOLOTL than by other methods (Fig. 5B-C). The advantages of AXOLOTL method are threefold in identifying abnormal gene expression of causal genes. Next, we further illustrate with detailed case examples.

**Figure 5.**
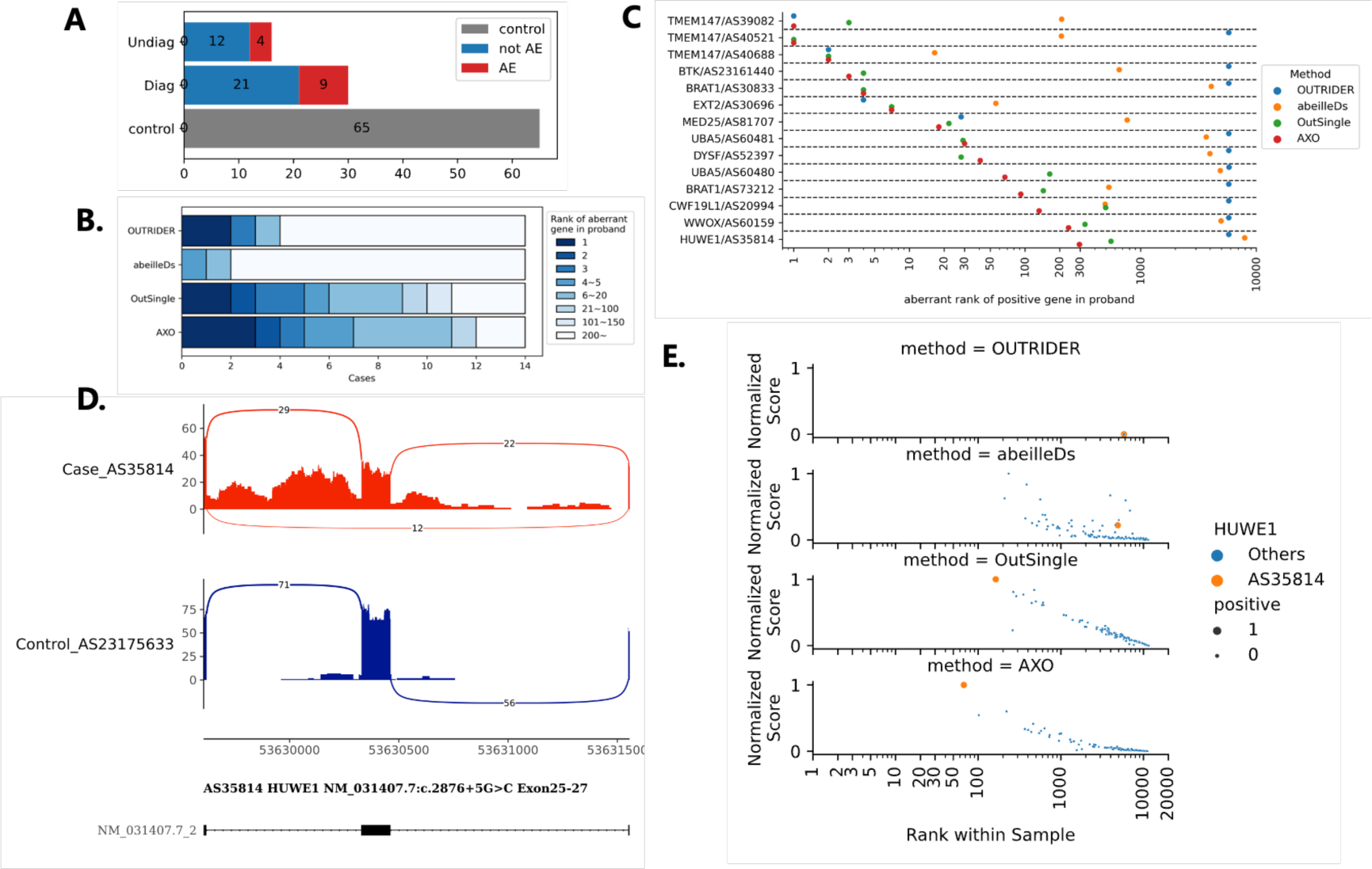
Performance comparison of AXOLOTL with other methods on blood RNA-seq samples of rare diseases dataset ly_111. **(A)** cohort structure and aberrant events identified by AXOLOTL in probands. **(B)** ranking distribution of aberrant gene in probands by different methods. **(C)** case-by-case comparisons of aberrant gene rankings in probands. **(D)** alternative splicing of HUWE1 gene in proband, compared to a representative control. **(E)** comparison of HUWE1 gene ranking and relative aberrant score by different methods. Horizontal axis of **(C)** and **(E)** represent log2 scaled of rankings.

First, AXOLOTL yields high rank for causal genes in diagnosed cases. In the male proband AS35814 (HP:0001263: global developmental delay; HP:0001274: agenesis of corpus callosum; HP:0001513: obesity) with dominant X-linked intellectual developmental disorder, the variant HUWE1 (chrX:53630324, c.2876+5G>C) caused alternative splicing via exon skipping (Fig. 5D). Aberrant ranking of the causal gene HUWE1 in the proband was 67 for AXOLOTL, which was higher than 163, 4890 and 5799 for OutSingle, ABEILLE and OUTRIDER, respectively (Fig. 5E). Similarly, the variant BTK (chrX:100613600:C>T, c.974+5G>A) of the male proband AS23161440 of recessive X-linked agammaglobulinemia (OMIM:300755) was ranked 18 by AXOLOTL, higher than 22, 763 and 28 by OutSingle, ABEILLE and OUTRIDER, respectively (Supplemental Fig. 2). In the male proband AS30696 (HP:0100777: Exostoses), who was diagnosed with the autosomal dominant variant EXT2 (chr11:44151690, c.1173+2T>A), EXT2 was ranked as the 4th most aberrant expressing gene by both AXOLOTL and OutSingle, compared with 5763 by OUTRIDER and 4072 by ABEILLE (Supplemental Fig. 2).

Second, AXOLOTL provides more reliable assessment on aberrant expression when sample duplicates of one proband or samples of multiple affected siblings are available. A female proband with BRAT1 variants (chr7:2581755:T>G, c.1014A>C; chr7:2583320:AG>A, c.706delC) have biological duplicate samples AS30833 and AS73212. The aberrant rankings of BRAT1 in two samples are 92;295 by AXO, while 144;554, 530;7934 and 5763;5764 by OutSingle, ABEILLE and OURIDER, respectively (Supplemental Fig. 3). In another example, the same compound heterozygous UBA5 variants (chr3:132394727:T>C, c.1091T>C; chr3:132384663:T>C, c.162-4T>C) were found in two siblings, AS60480 (1-year-old proband, HP:0001263) and AS60481 (7-year-old proband, HP:0001263 and HP:0001655: persistent foramen ovale). UBA5 was ranked higher in two probands by AXOLOTL than by other baseline methods (Supplemental Fig. 3).

Finally, AXOLOTL is more sensitive for detecting aberrantly expressed genes. Proband AS60159 (HP:0001250: seizure; HP:0002353: electroencephalogram abnormality) carries compound heterozygous WWOX variants (chr16:78191089:T>A, c.410-6991T>A; chr16:78198115:G>A, c.445G>A). The AS events at the junctions of interest were measured by the percent spliced in (PSI), which requires sufficient sequencing read coverage. Although the low abundance of WWOX mRNA in blood prevents AS detection of AS60159, AXOLOTL was among the top-3 aberrantly downregulated genes in the blood (Supplemental Fig. 4). In contrast, in this challenging case, WWOX was ranked as 4, 651 and 5770 by OutSingle, OURIDER and ABEILLE, respectively.

In another set of cases with aberrant expression, AXOLOTL and other methods had comparable performance in prioritizing causative genes (Table 1), including AS52397 with homozygous DYSF (chr2:71900503:C>T, c.5785-824C>T), ASD20994 with heterozygous CWF19L1 (chr10:102016016:C>A, c.504+3G>T) and three sibling probands, AS39082/AS40521/AS40688, with homozygous TMEM147 (chr19:36037846:G>T, c.345-1G>T). These results indicate that AXOLOTL can prioritize aberrant expression events that are less distinguishable via other methods. We concluded that AXOLOTL is an alternative method for detecting aberrant expression at high sensitivity.

## Conclusion and Discussion

Our comprehensive evaluations indicate that AXOLOTL provides a powerful approach for prioritizing aberrant gene expression. This study validates the effectiveness of AXOLOTL, as evidenced by its successful recovery of known cases from publicly available datasets as well as its resolution of challenging cases from our in-house dataset. In general, AXOLOTL consistently assigned higher rank to genes exhibiting aberrant expression in both the healthy and rare disease cohorts. It stands out by avoiding the use of arbitrary threshold, a common pitfall in other methods. Crucially, AXOLOTL exhibits robustness and sensitivity even in small cohorts, which makes it an attractive choice for a broad range of users.

The newly designed features of AXOLOTL are informative and have the potential to significantly enhance the accuracy of outlier detection. Normalization of gene expression levels addresses only technical confounders across samples and batches. It’s important to note that gene correlations are still not properly resolved by existing methods[7,13]. Considering this, we proposed the use of coexpression constraints as a measure of gene correlation. Based on this, we constructed sample-specific coexpression deviation features (*cts_dev*, *rank_dev*) derived from z score features (*ct*s_z, *rank_z*). AXOLOTL leverages sample-specific variability for the detection of gene aberrations. AXOLOTL is the first method that incorporates coexpression analysis into the detection of expression outliers. The ablation study and model explainability analysis demonstrated that the features provide complementary insights from various perspectives.

In the performance evaluation of AXOLOTL, artificial corruption was intentionally not employed. This approach contrasts with prior studies that incorporated read count foldchange corruption into random sample–gene pairs while leaving the remaining genes unchanged[7,13]. While the injection of artificial corruptions can be beneficial for method development purpose, we notice that fold-change corruption does not accurately represent any rare diseases. Real corruption of pathogenic genes results in downstream and feedback changes in the transcription of many genes. Given that aberrant gene expression events, as revealed by population-scale cross-tissue RNA-seq analysis, naturally occur in GTEx individuals [20], we think that the known aberrant events in GTEx are more real than artificially induced conditions. The performance evaluation of AXOLOTL on both publicly available rare disease datasets and GTEx datasets demonstrated the reliability of AXOLOTL in different CATs.

This study has several limitations. First, all methods demonstrated a low precision beyond a recall of 0.8, indicating that identifying a group of positive cases is challenging. In those cases, pathogenic genes with defects in splicing or protein function may not exhibit extreme gene expression levels. Second, AXOLOTL method is designed to improve the accuracy of detecting aberrancy in probands rather than evaluating pathogenicity. Nevertheless, the AXOLOTL score can be incorporated as RNA-seq-based supporting evidence in the prioritization of pathogenic genes, which is a hurdle in rare disease diagnosis. Third, this study proposed a coexpression constraint for the detection of aberrant gene expression. It would be worthwhile to investigate whether a similar correlation constraint of transcription events could aid in the detection of extreme alternative splicing and allele-specific expression.

In summary, our novel method improves the detection of aberrant gene expression, demonstrating its advantages in many real case studies. By highlighting the importance of coexpression as an essential biological constraint, we hope this work can ultimately empower data-driven diagnosis of rare diseases and inspire methodological innovations from new perspectives.

## Acknowledgments

We are grateful to the patients and their families for their participation in this research.

## Authors’ contributions

Conceptualization: WX. Supervision: WX, YW. Formal analysis: WX, FL, YL, YS, XL. Resources: YL, YW, JZ. Software: WX, FL Draft preparation: WX, FL, YL. All the authors read and approved the final manuscript.

## Data and code availability

The sequencing data of the ly111 dataset have not been deposited in a public repository to protect individual confidentiality. Our ethics approval and consent agreements allow us to share nonidentifiable patient data and gene expression count matrices data only. They are available for academic research in the Zenodo repository (https://zenodo.org/records/10408481). Supporting data of other datasets were downloaded from original publications. The source code of AXOLOTL method is available in GitHub (https://github.com/xuwenjian85/axolotl).

## Declarations

### Ethics, consent and permissions

This study was approved by the Institutional Review Board of the Capital Institute of Pediatrics (HERLLM2023060, SHERLLM2023061). Informed consent was obtained from all the subjects participating in the study. The research conformed to the principles of the Declaration of Helsinki.

### Consent for publication

We have obtained consent to publish from the participant to report individual patient data.

---

**Supplemental Figure S1.**
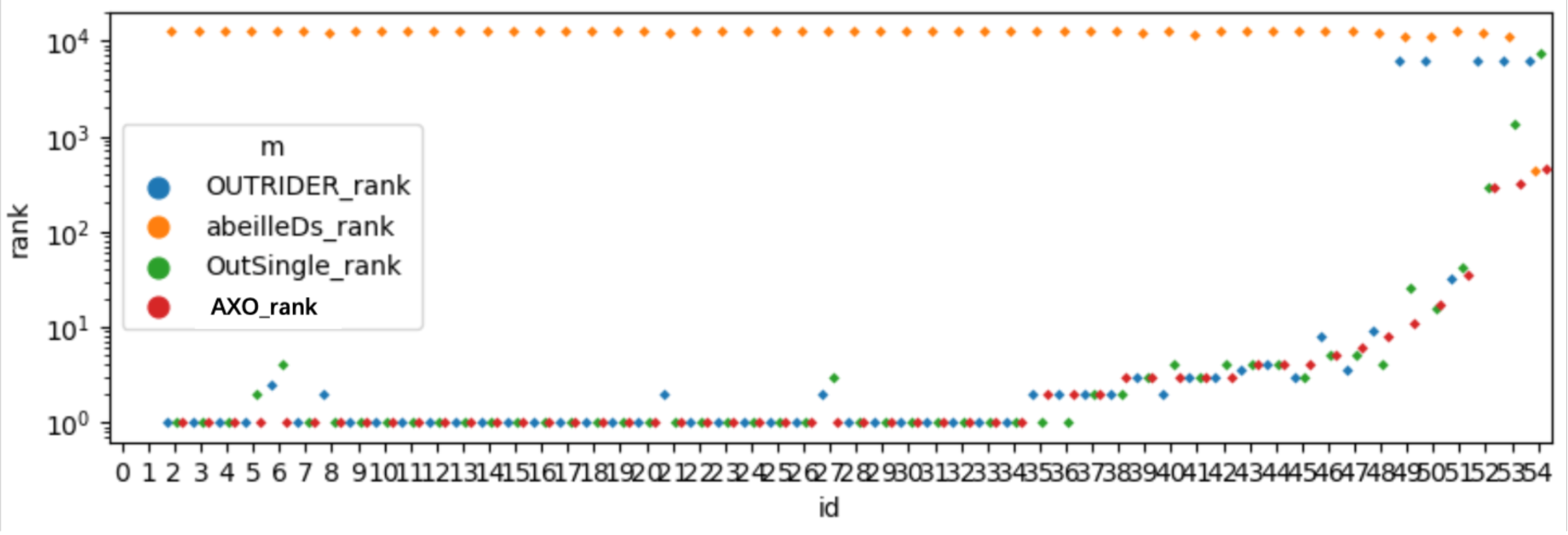
Performance comparison of AXOLOTL and other methods on pfib_423 dataset. The within-sample rankings of disease causative genes are shown for known positive samples.

**Supplemental Figure S2.**
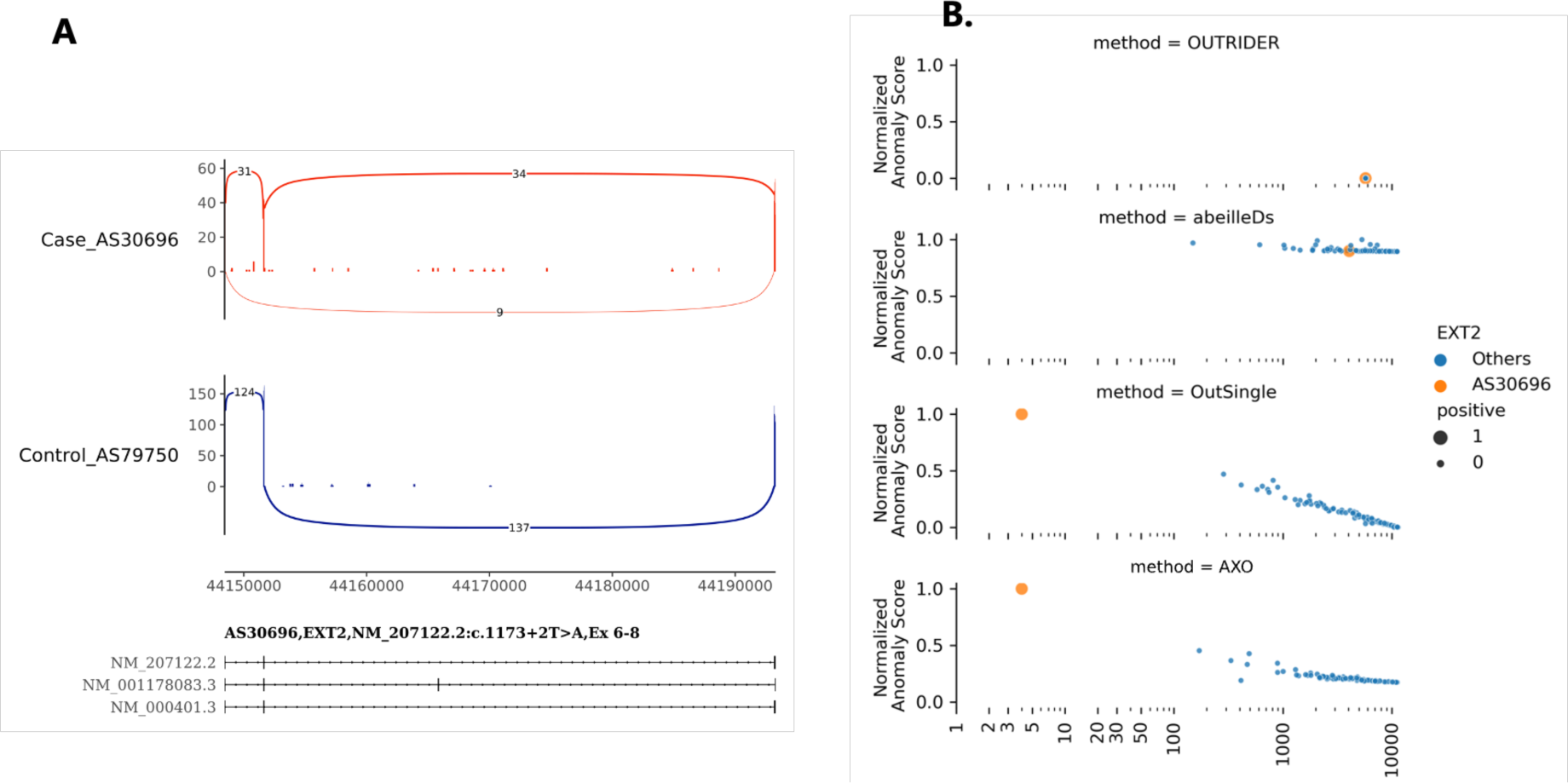
Performance comparison of AXOLOTL with other methods on dataset ly_111. **(A)** alternative splicing of EXT2 gene in proband, compared to representative control. **(B)** comparison of EXT2 gene ranking and relative aberrant score by different methods. Horizontal axis represents log2 scaled of rankings.

**Supplemental Figure S3.**
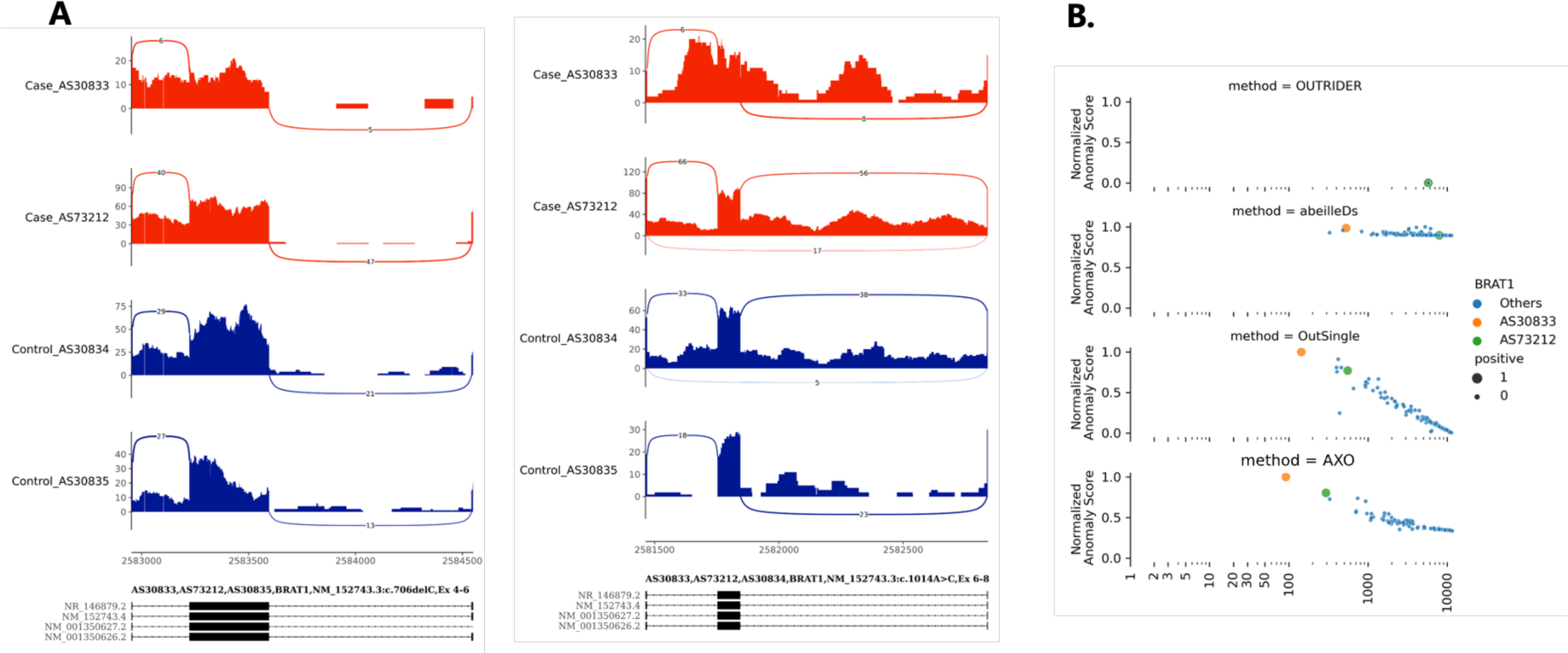
Performance comparison of AXOLOTL with other methods on dataset ly_111. **(A)** alternative splicing of BRAT1 gene in proband, compared to representative control. **(B)** comparison of BRAT1 gene ranking and relative aberrant score by different methods. Horizontal axis represents log2 scaled of rankings.

**Supplemental Figure S4.**
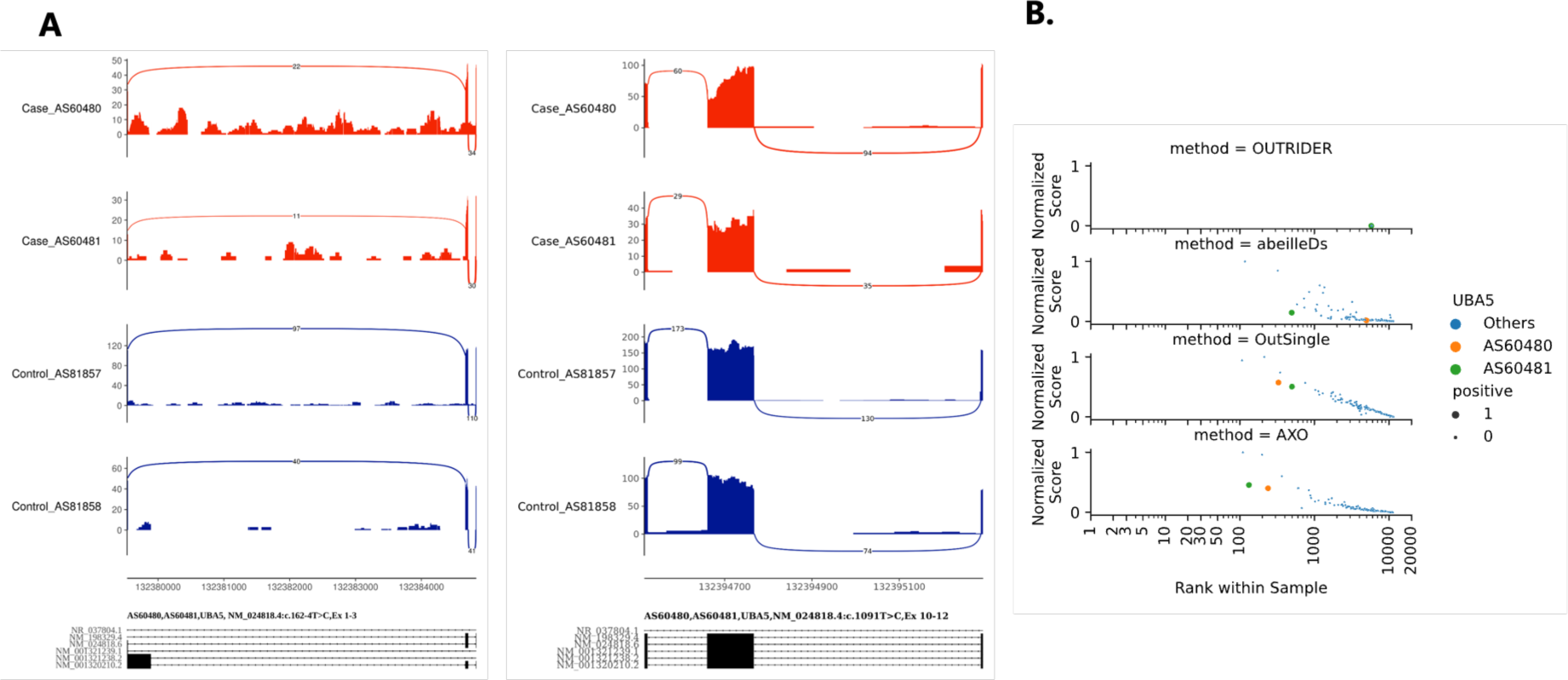
Performance comparison of AXOLOTL with other methods on dataset ly_111. **(A)** alternative splicing of UBA5 gene in proband, compared to representative control. **(B)** comparison of UBA5 gene ranking and relative aberrant score by different methods. Horizontal axis represents log2 scaled of rankings.

**Supplemental Figure S5.**
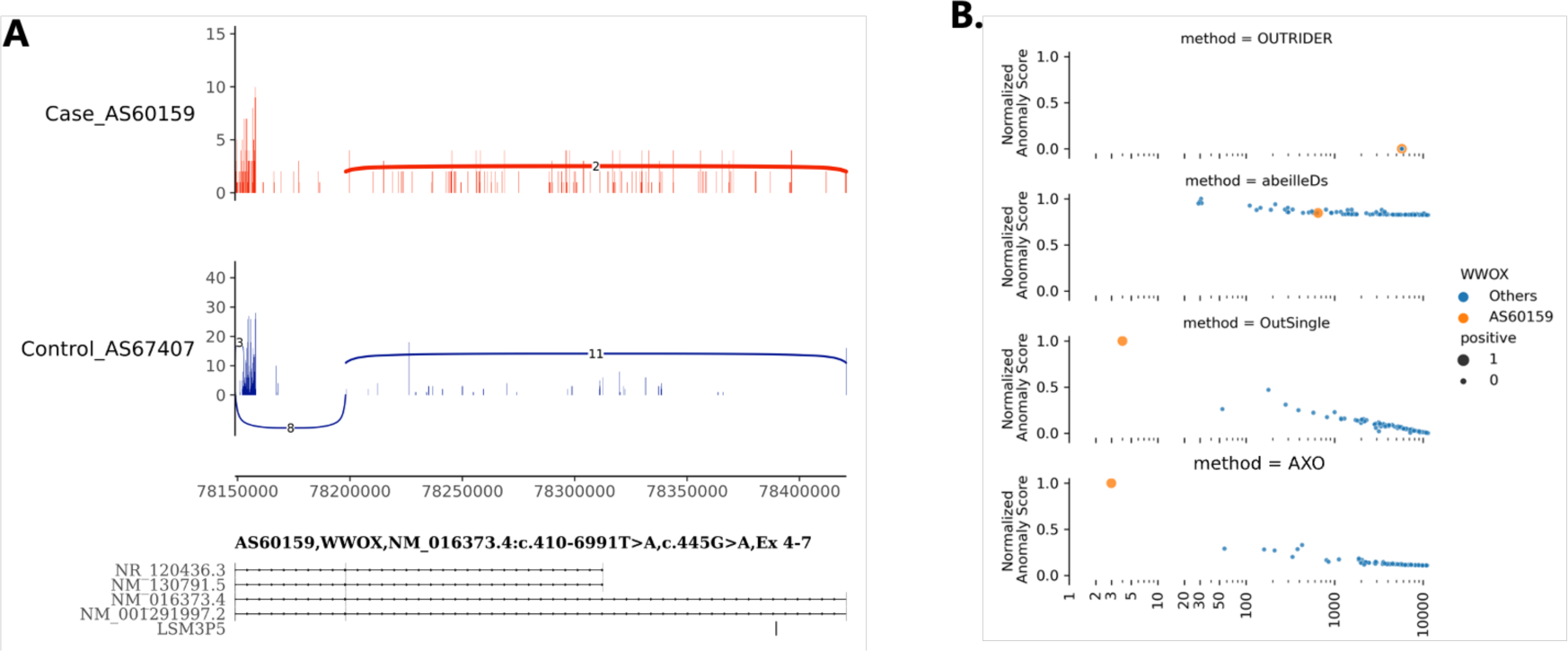
Performance comparison of AXOLOTL with other methods on dataset ly_111. **(A)** alternative splicing of WWOX gene in proband, compared to representative control. **(B)** comparison of WWOX gene ranking and relative aberrant score by different methods. Horizontal axis represents log2 scaled of rankings.

## Supplemental Methods

### Raw dataset preprocessing

#### **(1)** GTEx cohort - blood_755, skin_601, fib_504 and muscle_802

We used three files from GTEx Portal to define expression outliers in GTEx datasets. File1: GTEx_Analysis_v8_Annotations_SampleAttributesDS.txt

File2: GTEx_Analysis_2017-06-05_v8_RNASeQCv1.1.9_gene_reads.gct.gz File3: gtexV8.eOutlier.stats.globalOutliers.removed.txt.gz

##### Find high-quality samples of these tissues

File1 contains sample ids and sample annotations of all tissues. Samples of whole blood, skin - not sun exposed, cultured fibroblasts and muscle – skeletal were extracted into four GTEx datasets: blood_755, skin_601, fib_504 and muscle_802. We retrieved related samples using keywords ’*Whole Blood*’, ’*Skin - Not Sun Exposed (Suprapubic)*’, ’*Cells - Cultured fibroblasts*’ and ’*Muscle - Skeletal*’. We kept the high quality samples with SMRIN greater than 5.6. We use ’SAMPID’ and ‘SUBJID’ as sample ids.

##### Prepare read counts matrices

File2 are RNA-seq read counts (v8) provided by GTEx portal. Duplicate experiments of same samples are merged by keeping maximum count value. RNA-seq profiles of 4 tissues were separated as 4 data table. For each tissue, genes expressing all samples was kept. Multiple transcripts of the same gene were merged.

##### Define true expression outliers

File3 describes expression outliers in individual donors using z-score stat [1]. These outliers are globally aberrant expressing genes in multi-tissues, thus the outliers are supposed to be caused by special genetic background of healthy donors. We referred to the observed frequency of expression outlier in public rare disease datasets. Because overexpressing outliers are much more than under expressing outliers at the same Z-score amplitude, we define aberrant expression outliers as arbitrary cut-offs at Z-score >9 or <-7. The number of aberrantly expressed genes are, 36, 38, 43 and 24 for blood_755, skin_601, muscle_802 and fib_504 dataset. The outlier frequency is about 1 gene per 20 samples.

Expression matrices and true outliers are provided as **Supplemental Tables1-4.**

#### **(2)** Rare disease cohort 1 - pfib_423

Pfib_423, the largest publicly available rare disease RNA-seq cohort data to date, is a dataset of skin-derived fibroblasts of 423 patients (including samples from Kremer dataset[2]) suffering from mitochondrial disease or other Mendelian diseases [3]. Read count table was retrieved from Zenodo repository record 4646823 and 4646827. We compiled 59 positive aberrant expression events (**Supplemental Table 5**). The positives were reported in aberrant expression analysis and also confirmed by WES/WGS.

##### Define true expression outliers

Supplemental Tables of paper [3] (Table S2. Summary of cases diagnosed via RNA-seq, Table S3. Summary of candidate genes pinpointed via RNA-seq, and Table S4. Summary of WES-diagnosed cases with an RNA-defect) were manually merged. All aberrant expression events (labeled as ‘AE’ in the original publication) were extracted and used as 59 true outliers of the pfib_423 dataset.

##### Prepare read counts matrices

The RNA-seq read counts were downloaded as two parts, stranded specific table (fib_ss--hg19--gencode34/geneCounts.tsv.gz) and non-strand specific table (fib_ns--hg19--gencode34/geneCounts.tsv.gz). The two tables were merged. The genes with read count > 30 in over 95% of samples were kept for later analysis.

##### Simulation by subsampling pfib_423

Furthermore, *Pfib_423* is also used to analyze the robustness of AXOLOTL at settings of different sample size and different percent of positive patients. We set the sample size to [100, 150, 200, 250, 300] and the positive sample percentage to [0.05, 0.10, 0.15]. To obtain simulated datasets from pfib_423, we randomly sampled 100 times for each of 15 (sample-size, percentage) settings. 1500 simulated datasets were analyzed by AXOLOTL and baseline methods.

#### **(3)** Rare disease cohort 2 – pmuscle_36

Pmuscle_36 is a RNA-seq dataset of skeletal muscle biopsies of 22 COL6-RD patients and 14 age-matched controls [4]. COL6-RD is a form of congenital muscular dystrophy characterized by muscle weakness and joint defects. The molecular mechanism of COL6-RD is loss of function or dysfunction of COL6 microfibrils in the muscle extracellular matrix. COL6-RD can be caused by pathogenic variants in collagen VI genes (COL6A1, COL6A2, COL6A3). Pathogenic variants in COL6A genes were found in 22 patients. Immunostaining assays indicated that COL6A proteins were aberrantly low expressed in 5 patients and mis-localized in the other 17 patients. We defined COL6A genes of these 5 patients as positive expression outliers (**Supplemental Table 6**). Gene read counts were produced using our in-house pipeline described in [5] from RNA-seq raw data GSE103975.

#### **(4)** Rare disease cohort 3 – ly_111

##### Sample collection and variant pathogenicity assessment

We established stringent inclusion criteria for patients diagnosed with rare diseases. Firstly, the proband’s phenotype was utilized as a robust predictor of a monogenic rare disease.

Secondly, variants on suspected pathogenic genes were identified through WES/WGS, or Sanger sequencing results of the proband and parents, with annotation to the HGVS-nomenclature and classification according to the ACMG guideline. Thirdly, the proband presented uncertain significance (VUS), likely pathogenic (LP), or pathogenic (P) DNA variants with an alteration in splice effect (SpliceAI v1.3 score > 0.2) in the disease gene.

The investigation of variants of VUS necessitates the application of RNA-seq analysis to validate their predicted splicing defects. Our approach utilized blood RNA-seq on both resolved and unresolved samples. The pre- and post-RNA-seq splicing analysis classifications were delineated in columns ‘ACMG_DNA’ and ‘ACMG_RNAseq’ of **Table 1**. The former corresponds to the standard analysis with SpliceAI-predicted splicing effects, while the latter pertains to the variant classification based on WES/WGS and manually verified in Integrative Genomics Viewer (IGV). Splicing defects observed in RNA-seq data were considered as supporting evidence. The identification of alternative splicing events proximal to the VUS variant was instrumental in upgrading some variants to LP/P classification. Conversely, VUSs were maintained if no abnormal splicing findings were observed. Notably, through abnormal splicing analysis, we elevated the pathogenicity of selected VUS variants to LP/P and identified candidate genes in these cases.

##### Sample group definition

The classification of variants in the ’ACMG-RNAseq’ column represents the definitive basis for our conclusive diagnosis in each case: any likely pathogenic (LP) or pathogenic (P) variant is diagnosed, while VUS remains undiagnosed. Consequently, cases were categorized into diagnosed and undiagnosed groups based on this classification. Additionally, to expand the size of the ly_111 cohort, we incorporated additional samples to serve as a control group.

Diagnosed group (n=28). The samples have alternative splicing related LP/P level alleles, which are:

- 9 heterozygous cases affected by AD disorders,
- 5 homozygous cases affected by AR disorders,
- 12 compound heterozygous cases affected by AR disorders,
- 2 hemizygous males affected by X-linked disorders.

Undiagnosed group(n=18): the samples have only VUS level alleles, which are:

1. heterozygous cases affected by AD disorders,
2. heterozygous cases affected by AR disorders,
3. homozygous case affected by AR disorders,
4. compound heterozygous cases affected by AR disorders,
5. Heterozygous females affected by X-linked disorders

Control group 1: unaffected adult individuals in probands’ family (n=43). Control group 2: Miscellaneous samples (n=22):

probands affected by genes lowly expressed in blood;
probands who might have parental mosaic variants;
probands of copy number variant, covering a few exons of one gene or multiple genes.
probands and carriers of the thalassemia HBB variants. HBB gene is highly expressed in blood and is well-suited for the assessment of alternative splicing. However, the measurement of its expression level poses a challenge due to the considerable variability in HBB mRNA levels within blood samples.

##### RNA-seq, read alignment, quality control and splicing analysis

All samples were subjected to Whole Blood RNA-seq protocol and sequenced on Illumina NovaSeq 6000 platform with 150 bp paired-end reads. Total RNA was isolated with TRIzol-based RNA extraction. Quality of RNA was assessed by determination of the RNA integrity number (RIN) with a Bioanalyzer (Agilent). Next, polyadenylated RNA (mainly mRNA) was enriched with theNEBNext Poly(A) mRNA Magnetic Isolation Module (NEB, USA) followed by fragmentation, cDNA synthesis and library construction with the NEBNext Ultra™ II RNA Library Prep Kit for Illumina (NEB, USA). A minimum of 50 million reads were generated per sample. FASTQ files were processed with an inhouse pipeline. Raw reads were cleaned with fastp (v0.20.0), mapped with HiSat2 (v.2.1.0) to the GRCh38 GENCODE genome assembly and Gencode v41 as transcriptome reference. SNPs and indels were called using the GATK short variant discovery recommended pipeline with GATK (4.1.3.0) and STAR (2.5.3). Alternative splicing events were called by rMATS (v4.1.2) and visualized in the Integrated Genome Viewer (IGV; v2.16.2) and ggsashimi (v1.0.0).

##### True outlier labelling for AXOLOTL performance evaluation

Since our goal is evaluating the performance advantage of the AXOLOTL method, independent of the alternative splicing results, our focus centers on the validation of pathogenic genes through aberrant expression analysis. In this context, suspected pathogenic genes in probands are considered as true outliers. The aberrant rankings of these true outliers assigned by AXOLOTL and other methods is compared.

##### Implementation

The outlier detection models were implemented using the scikit-learn v0.21[6] with default parameters unless otherwise specified: Local Outlier Factor (LOF): *neighbors.LocalOutlierFactor (n_neighbors=20)*, Isolation Forest (IF): *ensemble.IsolationForest (n_estimators=20, max_samples =1.0)*, and One Class Support Vector Machine (OC-SVM): *svm.OneClassSVM (gamma=’auto’)*.

Pearson’s correlation coefficient (!) used to measure the strength of co-expression between genes was implemented by *scipy.stats*.*pearsonr* from scipy v1.3 [35].

Feature importance analysis of SHAP and LIME are implemented by scikit-explain v0.1.2.

Briefly, *skexplain.ExplainToolkit* was used to wrap the model LOF and feature matrix as *explainer* object. Then the SHAP and LIME results were retrieved from *explainer.local_attributions.* Next, by convert ‘shap_values LOF’ and ‘lime_values LOF’ values to importance scores for plotting purpose by *to_skexplain_importance*.

OUTRIDER v1.12.0 was installed on R-4.1.1 locally using R devtools command devtools::install_github (’gagneurlab/OUTRIDER’, dependencies=TRUE). Default parameters of OUTRIDER function is used.

ABEILLE v1.0.0 were implemented by R packages running on R-4.1.1. It is installed using R devtools command devtools::install_github(“UCA-MSI/ABEILLE”, dependencies=TRUE). The Nvidia V100-16GB GPU was utilized to accelerate the training of the tensorflow variational autoencoder model of ABEILLE.

## References

1. Kremer LS, Bader DM, Mertes C, Kopajtich R, Pichler G, Iuso A, et al. Genetic diagnosis of Mendelian disorders via RNA sequencing. Nat Commun. 2017;8:15824.

2. Cummings BB, Marshall JL, Tukiainen T, Lek M, Donkervoort S, Foley AR, et al. Improving genetic diagnosis in Mendelian disease with transcriptome sequencing. SCIENCE TRANSLATIONAL MEDICINE. 2017;12.

3. Yépez VA, Gusic M, Kopajtich R, Mertes C, Smith NH, Alston CL, et al. Clinical implementation of RNA sequencing for Mendelian disease diagnostics. Genome Med. 2022;14:38.

4. Frésard L, Smail C, Ferraro NM, Teran NA, Li X, Smith KS, et al. Identification of rare-disease genes using blood transcriptome sequencing and large control cohorts. Nat Med. 2019;25:911–9.

5. Murdock DR, Dai H, Burrage LC, Rosenfeld JA, Ketkar S, Müller MF, et al. Transcriptome-directed analysis for Mendelian disease diagnosis overcomes limitations of conventional genomic testing. J Clin Invest. 2021;131:141500.

6. Yepez VA, Mertes C, Muller MF, Klaproth-Andrade D, Wachutka L, Fresard L, et al. Detection of aberrant gene expression events in RNA sequencing data. Nature protocols. 2021;16:1276–96.

7. Brechtmann F, Mertes C, Matusevičiūtė A, Yépez VA, Avsec Ž, Herzog M, et al. OUTRIDER: A Statistical Method for Detecting Aberrantly Expressed Genes in RNA Sequencing Data. Am J Hum Genet. 2018;103:907–17.

8. Mertes C, Scheller IF, Yépez VA, Çelik MH, Liang Y, Kremer LS, et al. Detection of aberrant splicing events in RNA-seq data using FRASER. Nat Commun. 2021;12:529.

9. Jenkinson G, Li YI, Basu S, Cousin MA, Oliver GR, Klee EW. LeafCutterMD: an algorithm for outlier splicing detection in rare diseases. Bioinformatics. 2020;36:4609–15.

10. Dekker J, Schot R, Bongaerts M, De Valk WG, Van Veghel-Plandsoen MM, Monfils K, et al. Web- accessible application for identifying pathogenic transcripts with RNA-seq: Increased sensitivity in diagnosis of neurodevelopmental disorders. The American Journal of Human Genetics. 2023;110:251–72.

11. Love MI, Huber W, Anders S. Moderated estimation of fold change and dispersion for RNA-seq data with DESeq2. Genome Biology. 2014;15:550.

12. Salkovic E, Abbas MM, Belhaouari SB, Errafii K, Bensmail H. OutPyR: Bayesian inference for RNA-Seq outlier detection. Journal of Computational Science. 2020;47:101245.

13. Labory J, Le Bideau G, Pratella D, Yao J-E, Ait-El-Mkadem Saadi S, Bannwarth S, et al. ABEILLE: a novel method for ABerrant Expression Identification empLoying machine LEarning from RNA- sequencing data. Bioinformatics. 2022;38:4754–61.

14. Salkovic E, Sadeghi MA, Baggag A, Salem AGR, Bensmail H. OutSingle: a novel method of detecting and injecting outliers in RNA-Seq count data using the optimal hard threshold for singular values. Kendziorski C, editor. Bioinformatics. 2023;39:btad142.

15. Salkovic E, Bensmail H. A Novel Bayesian Outlier Score Based on the Negative Binomial Distribution for Detecting Aberrantly Expressed Genes in RNA-Seq Gene Expression Count Data. IEEE Access. 2021;9:75789–800.

16. Raina P, Guinea R, Chatsirisupachai K, Lopes I, Farooq Z, Guinea C, et al. GeneFriends: gene-co- expression databases and tools for humans and model organisms. Nucleic Acids Research. 2023;51:D145–58.

17. Langfelder P, Horvath S. WGCNA: an R package for weighted correlation network analysis. BMC Bioinformatics. 2008;9:559.

18. Saelens W, Cannoodt R, Saeys Y. A comprehensive evaluation of module detection methods for gene expression data. Nat Commun. 2018;9:1090.

19. GTEx Consortium. Genetic effects on gene expression across human tissues. Nature. 2017;550:204–13.

20. Ferraro NM, Strober BJ, Einson J, Abell NS, Aguet F, Barbeira AN, et al. Transcriptomic signatures across human tissues identify functional rare genetic variation. Science. 2020;369:eaaz5900.

21. Guadagnin E, Mohassel P, Johnson KR, Yang L, Santi M, Uapinyoying P, et al. Transcriptome analysis of collagen VI-related muscular dystrophy muscle biopsies. Ann Clin Transl Neurol. 2021;8:2184–98.

22. Xu W, He H, Guo Z, Li W. Evaluation of machine learning models on protein level inference from prioritized RNA features. Brief Bioinform. 2022;23:bbac091.

23. Lundberg SM, Lee S-I. A Unified Approach to Interpreting Model Predictions. In: Guyon I, Luxburg UV, Bengio S, Wallach H, Fergus R, Vishwanathan S, et al., editors. Advances in Neural Information Processing Systems [Internet]. Curran Associates, Inc.; 2017. Available from: https://proceedings.neurips.cc/paper_files/paper/2017/file/8a20a8621978632d76c43dfd28b67767-Paper.pdf

24. Ribeiro MT, Singh S, Guestrin C. “Why Should I Trust You?”: Explaining the Predictions of Any Classifier [Internet]. arXiv; 2016 [cited 2023 Jun 28]. Available from: http://arxiv.org/abs/1602.04938

25. Waskom ML. seaborn: statistical data visualization. Journal of Open Source Software. 2021;6:3021.

26. Robinson JT, Thorvaldsdóttir H, Winckler W, Guttman M, Lander ES, Getz G, et al. Integrative genomics viewer. Nat Biotechnol. 2011;29:24–6.

## Reference

1. Ferraro NM, Strober BJ, Einson J, et al. Transcriptomic signatures across human tissues identify functional rare genetic variation. Science 2020; 369:eaaz5900

2. Kremer LS, Bader DM, Mertes C, et al. Genetic diagnosis of Mendelian disorders via RNA sequencing. Nat. Commun. 2017; 8:15824

3. Yépez VA, Gusic M, Kopajtich R, et al. Clinical implementation of RNA sequencing for Mendelian disease diagnostics. Genome Med. 2022; 14:38

4. Guadagnin E, Mohassel P, Johnson KR, et al. Transcriptome analysis of collagen VI-related muscular dystrophy muscle biopsies. Ann. Clin. Transl. Neurol. 2021; 8:2184–2198

5. Xu W, He H, Guo Z, et al. Evaluation of machine learning models on protein level inference from prioritized RNA features. Brief. Bioinform. 2022; 23:bbac091

6. Pedregosa F, Varoquaux G, Gramfort A, et al. Scikit-learn: Machine Learning in Python. J. Mach. Learn. Res. 2011; 12:2825–2830

